# Identification of essential genes and fluconazole resistance genes in *Candida glabrata* by profiling of *Hermes* transposon insertions

**DOI:** 10.1101/2020.04.27.064048

**Authors:** Andrew N. Gale, Rima M. Sakhawala, Anton Levitan, Roded Sharan, Judith Berman, Winston Timp, Kyle W. Cunningham

**Affiliations:** Department of Biology, Johns Hopkins University, Baltimore, MD, USA; School of Molecular Microbiology and Biotechnology, Faculty of Life Sciences, Tel Aviv University, Tel Aviv, Israel; Blavatnik School of Computer Science, Tel Aviv University, Tel Aviv, Israel; Department of Biomedical Engineering, Johns Hopkins University, Baltimore, MD, USA

## Abstract

Within the budding yeasts, the opportunistic pathogen *Candida glabrata* and other members of the *Nakaseomyces* clade have developed virulence traits independently from the CTG clade that includes *Candida albicans*. To begin exploring the genetic basis of *C. glabrata* virulence and its innate resistance to antifungals, we launched the *Hermes* transposon from a plasmid and obtained more than 500,000 different semi-random insertions throughout the genome. Using machine learning, we identify up to 1278 protein-encoding genes (25% of total) that cannot tolerate transposon insertions and are thus essential for *C. glabrata* fitness in vitro. Interestingly, genes involved in mRNA splicing were less likely to be essential in *C. glabrata* than their orthologs in *S. cerevisiae*, whereas the opposite is true for genes involved in kinetochore function and chromosome segregation. Insertions in several known genes (e.g. *PDR1*, *CDR1*, *PDR16*, *PDR17*, *UPC2A*, *DAP1*) caused hypersensitivity to the first-line antifungal fluconazole, and we identify 12 additional genes that also contribute to innate fluconazole resistance (*KGD1*, *KGD2*, *YHR045W*, etc). Insertions in 200 other genes conferred significant resistance to fluconazole, two-thirds of which function in mitochondria and likely down-regulate Pdr1 expression or function. These findings show the utility of transposon insertion profiling in genome-wide forward-genetic investigations of fungal pathogens.

**IMPORTANCE:** Pathogenic yeasts cause mucosal and systemic infections in millions of people each year. The innate resistance of *Candida glabrata* to fluconazole and its ability to acquire resistance to 2 other antifungals are contributing to its rise in incidence. Our understanding of *C. glabrata* biology has been hampered by inefficient genetic and genomic tools. This study addresses those deficiencies by developing powerful transposon mutagenesis strategies for the first time in this pathogen. We identify nearly all essential genes of *C. glabrata* that could be targeted for development of new antifungals. We generate large pools of random insertion mutants that can be easily monitored *en masse* with deep sequencing, thus enabling identification of genes involved in any number of biological processes. We identify dozens of new genes that increase or decrease innate resistance of clinical isolate BG2 to fluconazole and provide resources for further exploration of *C. glabrata* genetics and genomics.

## INTRODUCTION

The yeast *Candida glabrata* is the second most common cause of candidiasis in humans and its incidence is rising due to its innate resistance to the first-line antifungal fluconazole (1, 2). The species acquires further resistance to fluconazole and related azole-class antifungals through mutations that overexpress the drug target (Erg11) and increase drug efflux (e.g. Pdr1) (3, 4). Though *C. glabrata* is naturally haploid, chromosomal aneuploidies and rearrangements can also contribute to antifungal resistance and to persistent infections (5, 6). The species is closely related to pathogenic *C. bracarensis* and *C. nivariensis* species and non-pathogenic members of the *Nakaseomyces* clade such as *N. delphensis* and *N. bacillisporus* (7). This group is also more closely related to the non-pathogenic bakers’ yeast *Saccharomyces cerevisiae* than to *Candida albicans* and other pathogenic yeasts of the CTG clade (8). This evolutionary relationship raises many questions of how pathogenicity, virulence, host colonization, drug resistance, and drug tolerance have evolved independently among budding yeasts.

Conventional genetic and genomic approaches are much more arduous in *C. glabrata* than *S. cerevisiae* because it lacks a sexual cycle and gene knockouts and knockins remain inefficient, even when aided by CRISPR/Cas9 technology (9, 10). Previously, a consortium of researchers has replaced ~15% of the non-essential genes with DNA barcodes and a nourseothricin-resistance marker (*NATr*) (11). Systematic screens of this collection have yielded insights into a number of biological processes including relative fitness, colony morphology, biofilm formation, and resistance to three different classes of clinical antifungals (fluconazole, caspofungin, amphotericin B). However, the collection is highly biased toward regulatory genes of interest to the community. In a pioneering approach, thousands of random *Tn7* transposon insertions also were generated *in vitro* and integrated into the *C. glabrata* genome, which were then arrayed and screened individually for susceptibility to antifungals (12, 13).

Transposon insertion profiling in eukaryotes has dramatically improved in recent years. Next-generation DNA sequencing technology facilitates *en masse* analyses of very large pools of insertion mutants (14–16). Typically, genomic DNA adjacent to each insertion site is directly amplified by PCR and sequenced using Illumina technology, and then the reads are mapped to precise sites in the genome and tabulated. In contrast to CRISPR/Cas9 screening approaches that are user-guided and indirect, transposon insertion profiling enables direct observation and quantitation of each mutation in the population and full coverage of the genome. In *S. cerevisiae*, millions of independent insertions have been obtained using derivatives of the maize *Activator/Dissociation* (*mini*-*Ac/Ds*) transposon (16) and the housefly *Hermes* transposon (17). In *C. albicans*, a *mini-Ac/Ds* transposon (18) and a *PiggyBac* transposon (19) have yielded hundreds of thousands of defined insertions. In the fission yeast *Schizosaccharomyces pombe*, which is distantly related to all the budding yeasts, both *Hermes* (14) and *PiggyBac* (20) have produced high-density insertion pools. Remarkably, all the transposons inserted with high enough density to reliably distinguish most non-essential genes from essential genes, where severe fitness defects of insertion mutants cause their depletion from the pool (21).

In this study, we adapt the *Hermes* transposon for insertion profiling in a clinical isolate of *C. glabrata*. Over 500,000 different insertion sites were identified using the QIseq method (15), allowing for the first description of its essentialome and comparison with other species. We show that *Hermes* inserts in *C. glabrata* with sequence bias and nucleosome bias similar to *S. cerevisiae* but with a much stronger centromere bias, suggesting a difference in chromosome architecture. Interestingly, genes involved in kinetochore function were more likely to be essential in *C. glabrata* than *S. cerevisiae*. We also identify hundreds of genes that alter susceptibility to fluconazole, including many previously identified genes. These findings illustrate the power of *in vivo* transposon mutagenesis when coupled with next-generation DNA sequencing for functional genomics research. They also extend our understanding of drug resistance mechanisms that operate in an important pathogen of humans, thus facilitating development of improved and novel antifungal strategies.

## RESULTS

### *Hermes* transposition in *C. glabrata*

Our strategy for launching the *Hermes-NATr* transposon and enriching for insertions in *C. glabrata* was based on prior studies in *S. cerevisiae* (17) and *S. pombe* (14). Briefly, a centromere-containing plasmid bearing a methionine-repressible *MET3* promoter and a counter-selectable *URA3* gene (22) was modified to express the *Hermes* transposase and to contain a *Hermes* transposon in which a *NATr* expression cassette replaced the natural transposase gene. Upon induction of transposase expression, the transposon can be excised from the plasmid and inserted into the genome. The plasmid launchpad is frequently lost and the cells become resistant to 5-FOA while remaining resistant to nourseothricin. After transformation of *C. glabrata* strain BG14 (23) and growth in synthetic complete medium lacking uracil, we observed a much greater number of insertion mutants (simultaneously resistant to both 5-FOA and nourseothricin) in the absence of methionine and cysteine than in their presence. The number of insertion mutants increased 50-fold or more after the cultures reached stationary phase on day 1 (Supplemental Fig. S1). Thus, after 3 days of induction, the vast majority of insertion mutants are likely to be independent of one another with relatively few insertions that occurred during growth phase and proliferated to a disproportionate frequency (i.e. jackpots).

In three separate cultures, less than 0.04% of cells in the population acquired an insertion into the genome, suggesting that instances of re-excision and re-insertion in the same cell line are extremely rare. Insertion mutants were enriched to high frequency (>99%) by passaging several times in medium containing nourseothricin and 5-FOA (see Materials and Methods). The enriched pools were frozen in aliquots for future use. Genomic DNA was extracted from each pool, sheared, repaired, ligated to adaptors, and amplified by PCR with custom primer pairs that target the adaptor and the right inverted repeat of the *Hermes* transposon as described previously (15, 21). The resulting libraries of PCR products were then directly sequenced on an Illumina MiSeq instrument using a custom primer that hybridized to the end of the transposon. Sequence reads were filtered for quality and mapped to the reference genome of *C. glabrata* strain CBS138 (see Materials and Methods). Sequence reads that map to the same insertion site were tallied and visualized using the IGV genome browser together with annotated genes and other genomic features (Fig. 1). The results appeared comparable to previous findings in *S. cerevisiae*, where insertions are distributed semi-randomly throughout the lengths of all 13 chromosomes of *C. glabrata*. Overall, the three pools contained between 164,000 and 335,000 defined insertions, with a combined total of 513,123 unique insertion sites. Approximately 4% of the CBS138 genome was unmappable using short reads of BG14 genomic DNA and another 17% contains essential genes (see below). Excluding these regions, an average of one insertion was observed every 19 bp.

**Figure 1.**
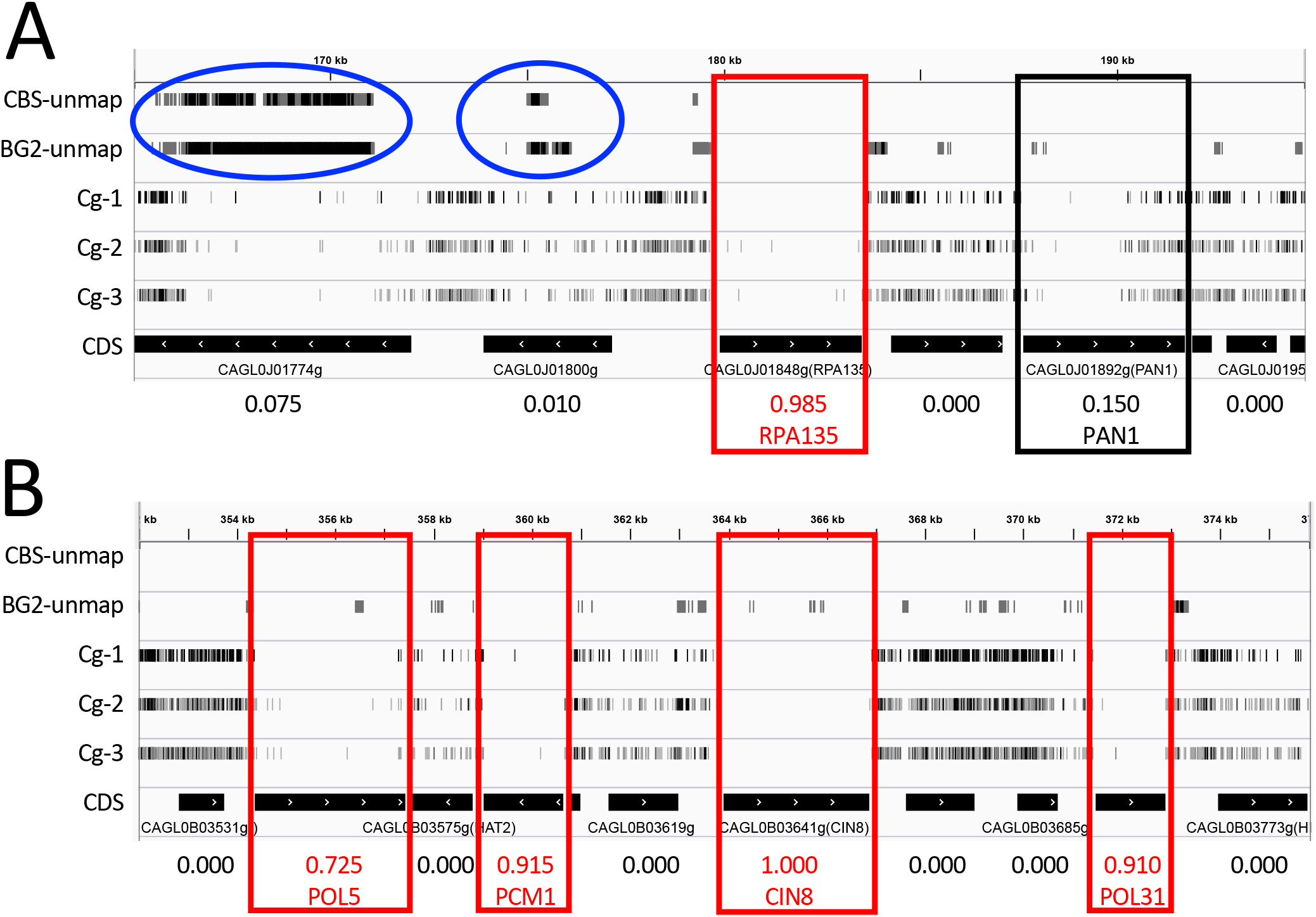
*Hermes-NATr* insertions visualized in *C. glabrata*. IGV browser representations of insertions in segments of chromosomes J (A) and B (B). Each row contains tick marks representing mapped insertion sites within a particular sequenced library that have been scaled to reflect read counts at that site. Segments of the CBS138 reference genome that are unmappable with short reads of CBS138 and BG2 genomic DNA are depicted in the top two tracks and highlighted (blue circles). Black bars indicate the positions of coding sequences and arrows indicated direction of transcription. Numbers at bottom indicate essentiality scores. Essential genes (red boxes) and *PAN1* (black box) are indicated.

### Insertion biases

*Hermes* transposons are known to insert preferentially at sites with T at position +2 and A at position +7 (17). In the three *C. glabrata* pools of *Hermes-NATr* insertions, 72% of all mapped sequence reads were located at such TA sites, a frequency that is 7.8-fold higher than that of TA sites in the genome (Fig. 2A). Near-cognate TG and CA sites exhibited 1.15- and 1.12-fold enrichment, while the nine non-cognate (non-T and non-A) sites exhibited 10- to 162-fold underrepresentation (Fig. 2A). In spite of this 1,250-fold range of insertion site bias, all annotated genes of *C. glabrata* contain multiple preferred TA sites and near-cognate sites. These findings suggest that with sufficient diversity of insertions in the initial pool, complexity of the library, and depth of sequencing, all genes can be profiled.

**Figure 2.**
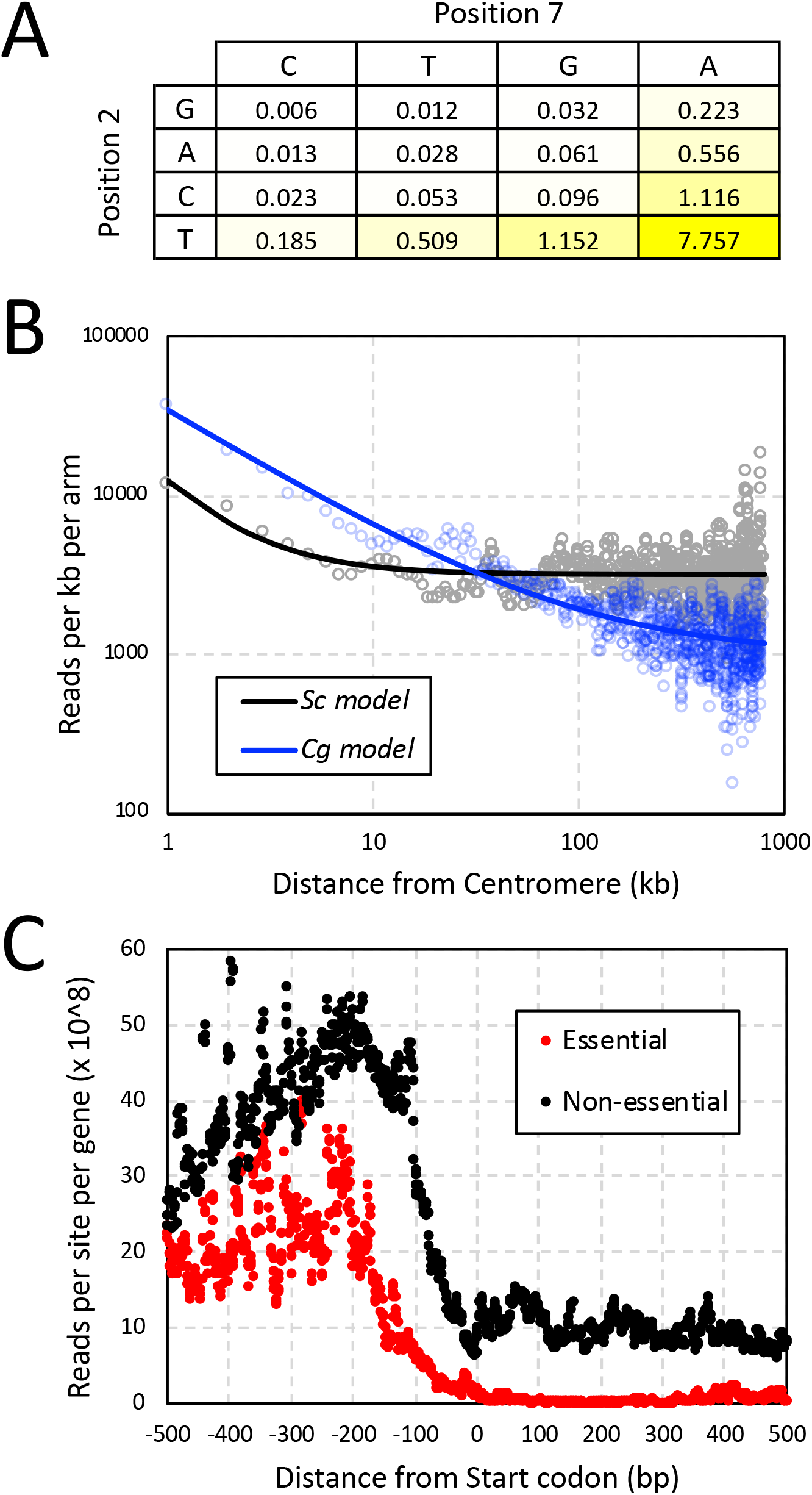
Factors biasing the sites of *Hermes-NATr* insertions. (A) The preferences for specific nucleotides at positions 2 and 7 were calculated by dividing the frequency of sequence reads at each site (obtained from libraries derived from three independent pools) by the frequency of such sites in the *C. glabrata* genome. (B) The number of sequencing reads within 1 kb bins were tabulated across all chromosome arms beginning at the centromeres. Smooth lines indicate non-linear regression to a standard power function. (C) The number of sequencing reads at each nucleotide position relative to the start codon were tabulated for all 1-to-1 non-essential genes (black) and essential genes (red) and divided by the number of genes in each set.

*Hermes* and other transposons also insert more frequently into DNA that is physically close to the site of their excision. As *Hermes-NATr* was launched from a centromere-containing plasmid and centromeres tend to cluster within the nucleus of *C. glabrata* (24), we expect to observe an insertion bias toward all centromeres. To quantify the magnitude of this bias, the read counts along all 26 chromosome arms of *C. glabrata* were tabulated in 1 kb increments from the centromere and compared to data from 32 chromosome arms of *S. cerevisiae* (21). Though insertions were biased toward centromeres in both species, the effect appeared much stronger and longer in *C. glabrata* relative to *S. cerevisiae* (Fig. 2B). This difference suggests potential variation between the species in architecture and/or packaging of chromosomes in the nucleus.

Transposon insertions *in vivo* are also biased toward nucleosome-free regions (17), such as promoter regions upstream of coding sequences. By aligning all non-essential genes at the start codons and tabulating read counts at every nearby position, a clear bias toward the 5’ non-coding region was observed relative to coding sequences (Fig. 2C, black). Within the coding sequences of non-essential genes, read counts oscillated slightly with a periodicity of approximately 170 bp, due to the positioning of nucleosomes in many genes (25). Essential genes, as defined by a machine learning approach in the next section, were depleted of read counts in the coding sequences by > 30-fold (Fig. 2C, blue) and also were depleted significantly in the adjacent 5’ non-coding region between −200 and −1 relative to the start codon (Fig. 2C, red). These findings suggest *Hermes-NATr* insertions in *C. glabrata* have little or no enhancer and promoter activities, which is similar to *Hermes-NATr* in *S. cerevisiae* (21) and distinct from *mini-Ac/Ds* that contains such activities (16).

### Identification of essential genes

Essential genes perform functions that are critical for growth and survival in the laboratory and their identification can facilitate development of new antifungals. Generally, transposon insertions within the coding sequences of essential genes greatly diminish competitive fitness, leading to depletion of sequence reads relative to those in the surrounding non-essential DNA. To discover essential genes in *C. glabrata*, we combined the data from all three pools after normalizing for the 4-fold under-sequencing of libraries from pools Cg-2 and Cg-3 (see Materials and Methods) and then we implemented a machine learning approach that had been applied successfully to multiple transposon insertion datasets from multiple species (21). The algorithm was trained on a set of confirmed essential and non-essential protein-encoding genes from *S. cerevisiae* using eight classification features that represent different aspects of essentiality. An essentiality score ranging from 0 (non-essential) to 1 (essential) was calculated for each gene. A histogram of the output revealed a bimodal distribution with maxima in the first and last deciles and relatively few genes in the middle quartiles (Fig. 3A). A total of 1342 out of 5282 protein-encoding genes scored above 0.5 and were classified as essential for competitive fitness in these conditions (Supplemental Table S1). This number dropped to 1278 out of 5190 genes (24.6%) after filtering genes that contain > 50% unmappable segments, which is similar to 1232 out of 5674 genes (21.7%) in *S. cerevisiae* cultivated under similar conditions (21). Hundreds of genes in *C. glabrata* and *S. cerevisiae* exist as duplications from an ancient whole genome duplication (WGD) event while thousands exist as singletons (26). When only 1-to-1 orthologous genes are considered (3880 genes total) (27), 84% of essential genes in *C. glabrata* were also essential in *S. cerevisiae*.

**Figure 3.**
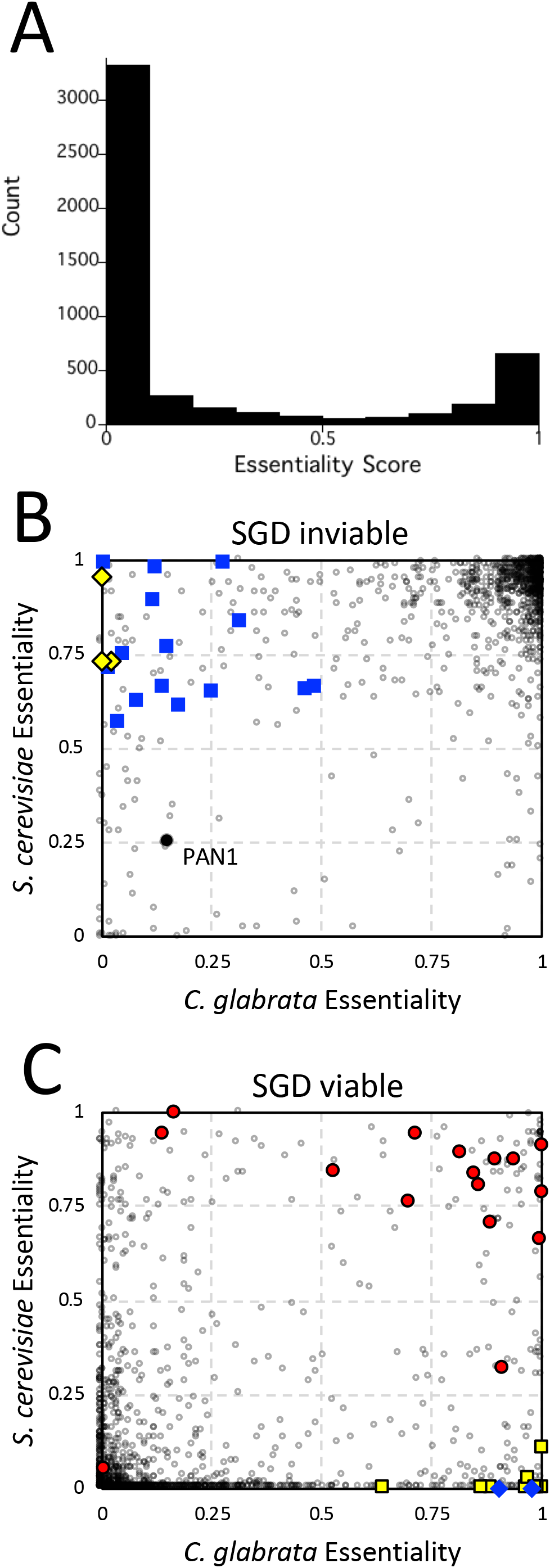
Essentiality comparisons between species. (A) Histogram of essentiality scores for all protein-encoding genes of *C. glabrata*. The 1-to-1 orthology group was split into two groups based on SGD annotations for inviable (B) and viable (C). Dark gray circles indicate individual othologous genes. Spliceosomal complex genes (blue squares) and SPS genes (yellow diamonds) were scored as essential in *S. cerevisiae* but not *C. glabrata*. Spermidine biosynthesis genes (blue diamonds) and kinetochore complex genes (yellow squares) were scored as essential in *C. glabrata* but not *S. cerevisiae*. V-ATPase genes (red circles) and *PAN1* (black circle) are indicated.

To explore possible instances of essentialome evolvability, we focused on 1001 genes of the 1-to-1 orthology group that were confirmed as inviable (essential) in *S. cerevisiae* through conventional approaches (28). We found 93 genes of *C. glabrata* that scored below 0.5 (non-essential) while scoring 0.5 or higher in *S. cerevisiae* (essential) and annotated as inviable at the Saccharomyces Genome Database (Fig. 3B). Gene ontology (GO) analyses (29) revealed enrichment of spliceosomal complex (GO:0005681, P-value = 1.8E-4, FDR q-value = 1.2E-1) encompassing 15 genes (Fig. 3B, blue squares) and transcriptional mediator complex (GO:0016592, P-value = 8.6E-4, FDR q-value = 2.9E-1) encompassing 5 genes. *C. glabrata* retains less than half as many introns as *S. cerevisiae* (30), so perhaps the level of importance of some spliceosomal subunits has been relaxed in this species. Another interesting example is the SPS complex, where all 3 components (*SSY1*, *PTR3*, *SSY5*) were essential in *S. cerevisiae* but not *C. glabrata* (Fig. 3B, yellow diamonds). The SPS complex induces expression of amino acid transporters and are only essential in auxotrophic strains of *S. cerevisiae* (31) such as the BY4741 strain (*his3Δ*, *leu2Δ*, *met15Δ*) and are non-essential in the prototrophic BG14 strain of *C. glabrata*. Last, the ortholog of *S. cerevisiae VHT1* in *C. glabrata* (CAGL0K04565g) scored as non-essential when it is critical for uptake of the essential vitamin H (biotin) present in the medium. Surprisingly, BLAST searches revealed a duplicate of CAGL0K04565g termed CAGL0K04609g nearby on chromosome K and this duplicate gene scored as essential. Phylogenetic analyses of these genes and their orthologs from other species indicate that the duplication is preserved in one sub-clade of the Nakaseomyces (*C. glabrata, C. bracarensis, C. nivariensis, N. delphensis*) but not in the other sub-clade (*C. castellii, N. bacillisporus*) or in any other species of yeasts (not shown). In the four species with duplicates, the ancestral CAGL0K04565g orthologs evolved more than twice as fast as the duplicated CAGL0K04609g orthologs (Supplemental Figure 2), suggesting that the duplicates are more likely to retain the essential function in biotin transport. Recently, individual knockouts of the CAGL0K04565g and CAGL0K04609g genes in *C. glabrata* showed that only the latter was essential for growth in these conditions (32), thus validating our assessments with transposon mutagenesis.

On the other end of the spectrum, a total of 138 genes were scored as essential in *C. glabrata* and not *S. cerevisiae* where they also have been confirmed as non-essential. GO term analyses suggest polyamine biosynthesis (GO:0006596, P-value 5.23E-6, FDR 2.29E-2) involving the *SPE1*, *SPE2*, and *SPE3* genes is one such crucial process (Fig. 3C, blue diamonds). Mutations in these genes in *S. cerevisiae* cause spermidine auxotrophy, but unlike the *C. glabrata* insertion mutants the *S. cerevisiae* insertion mutants exhibit only mild fitness defects in this spermidine-free culture media. Another major GO term component enriched in this set was defined by eight genes that contribute to the kinetochore (GO:0000776, P-value = 5.76E-5, FDR q-value = 6.39E-3; Fig. 3C, yellow squares). Essentiality of one such kinetochore gene (*CIN8*) can be explained by the loss of the functionally redundant *KIP1* gene in *C. glabrata*. In *S. cerevisiae*, *CIN8* non-essential in wild-type strains essential in *kip1Δ* strains (33). Similarly, the essentiality of several other kinetochore genes in this GO category (*BUB1, BUB2, BUB3, CBF1*, *CTF19*, *MCM21*, *MCM22*, *KAR3* as well as *CIN8*) are potentially explained by the absence of *BIR1. BIR1* (CAGL0M09152g) is currently annotated as a pseudogenein the *C. glabrata* reference genome (8) because of an in-frame stop codon that occurs at position 588, which would eliminate the strongly conserved C-terminal domain that is essential for chromosome segregation in *S. cerevisiae* (34) and mammalian cells (35). In *S. cerevisiae*, *bir1* hypomorphic mutations exhibit synthetic lethality with all these kinetochore genes (34, 36). However, *BIR1* is not likely to be a pseudogene in *C. glabrata* because the C-terminal domain is well-conserved in the +1 reading frame, a single mRNA is expressed that spans the entire gene (37), the mRNA contains a 7 base pair programmed +1 ribosomal frameshift sequence (38) just upstream of the in-frame stop codon, and all of these features are conserved in all five sequenced species of the *Nakaseomyces* clade (Supplemental Fig. S3). If full-length Bir1 is expressed through a programmed +1 ribosomal frameshifting mechanism in these species, this mechanism could contribute to chromosome instability, which previously has been associated with acquired fluconazole resistance (5) and persistent infections (6).

Some duplicates from the WGD event have been fully retained in both species, forming the 2-to-2 orthology group with 606 genes. Another 236 singleton genes of *C. glabrata* remain duplicated in *S. cerevisiae* while 182 duplicated genes in *C. glabrata* are singletons in *S. cerevisiae*, forming the 1-to-2 and 2-to-1 orthology groups. For the singletons of both species, the frequency of essentiality (30%) was similar to that of the 1-to-1 group (29%). But when a duplicate exists, the frequency of essentiality is much lower (7.5%), suggesting that duplicated genes in both species often function redundantly in essential processes. Thus, essentiality of the singleton in the 1-to-2 and 2-to-1 orthology groups can help expose redundant essentiality in the species with non-essential duplicates. In one example, the singleton *AUR1* gene in *S. cerevisiae* is essential while neither of the duplicates in *C. glabrata* is essential, though both species remain highly susceptible to the compound aureobasidin A that directly inhibits the products of *AUR1* genes (39).

The 1-to-0 and 0-to-1 orthology groups consisted of 33 and 126 genes, respectively, after excluding *BIR1* and all genes that were not present in the common ancestor of both species. Of these, only one gene in *C. glabrata* (CAGL0M04543g) and only three genes in *S. cerevisiae* scored over 0.5. All four are likely to be false positives due to their small size (228 - 540 bp) or borderline score. False positives also can occur when genes have large segments of repetitive DNA in which insertions cannot be mapped accurately with short-read Illumina technology and were therefore filtered out of our datasets. We find 1.4% of the CBS138 reference genome is unmappable even with perfect 75 bp matches to the reference genome and 4.4% is unmappable when BG14 DNA is used. The number of unmappable sites within each gene was tabulated to facilitate calling of essential genes (Supplemental Table S1). Out of 82 non-essential genes in the 1-to-1 orthology group that scored as essential in both species, four genes involved in mating type determination were almost completely unmappable and thus erroneously scored as essential when the insertions in these genes were simply filtered from the set of mapped reads. Interestingly, dozens of other genes scored as essential in both *C. glabrata* and *S. cerevisiae* have been validated as non-essential in *S. cerevisiae* and lack unmappable segments. Twelve of these mutants encode critical subunits of the V-ATPase (Fig. 3C, red circles), suggesting this enzyme complex strongly contributes to fitness in both yeasts in the conditions used to generate the pools. Note that some essential genes score as non-essential in both species, often because the gene products contain large non-essential domains at their C-termini adjacent to essential N-terminal domains (e.g. *PAN1* in Fig. 1A, and Fig. 3B, black circle). Altogether, transposon insertion profiling and analysis by machine learning can generate interesting insights into genes and processes that are essential for fitness, though care must be exercised to identify false positives and negatives.

### Genes that regulate susceptibility to fluconazole

The large pools of *Hermes* insertion mutants and the profiling methods described above provide an unprecedented opportunity to identify genes that critically impact fitness of *C. glabrata* in new conditions. To identify genes that regulate susceptibility of *C. glabrata* to fluconazole, we grew pool Cg-1 to stationary phase, diluted aliquots 100-fold into fresh medium containing or lacking fluconazole (128 μg/mL), and shook the planktonic cultures for 24 hr at 30°C. Cells were then pelleted, washed free of drugs, and cultured in an equal volume of fresh medium for an additional 48 hr. These conditions were chosen to expose genes that alter cell survival and recovery rates, in addition to genes that alter sensitivity and resistance to the antifungal. Genomic DNA from the saturated cultures was then isolated, converted into sequencing libraries, and sequenced as before. For each annotated gene, the total number of reads that mapped within the coding sequences were tabulated (Supplemental Table S2). Each gene was then represented as a single point in a 2-dimensional plot of experimental (fluconazole) versus control conditions (Fig. 4A). Most genes appear very close to the main diagonal and therefore do not play a strong role in susceptibility to fluconazole. Genes below the main diagonal promote resistance (i.e. confer hypersensitivity when disrupted with transposons) to fluconazole whereas genes above the main diagonal promote sensitivity to fluconazole. Significance of each gene was calculated as z-score relative to the local standard deviation (see Materials and Methods).

**Figure 4.**
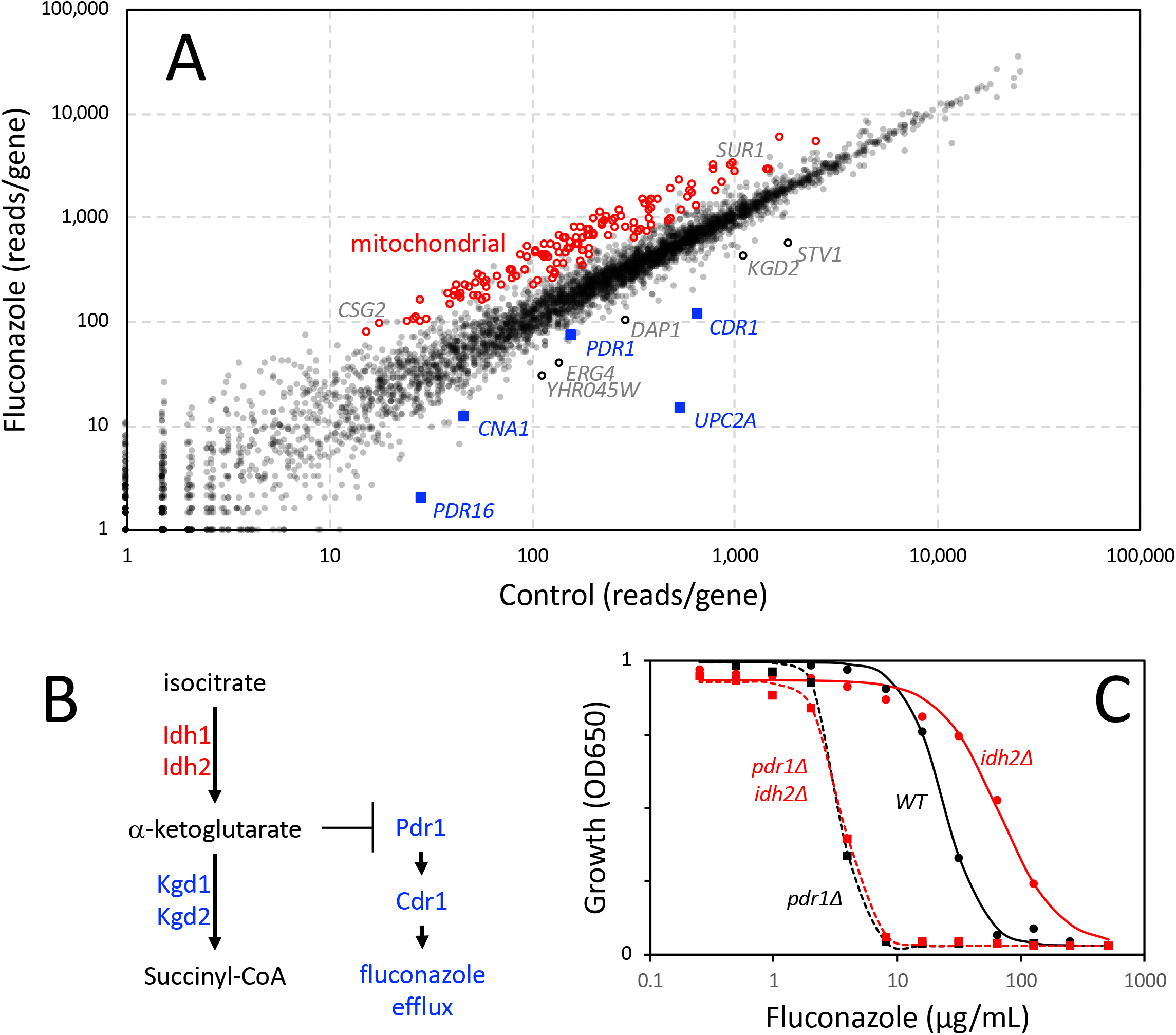
Identification of fluconazole susceptibility genes in *C. glabrata*. (A) Pool Cg-1 was split into 4 portions and duplicates were regrown in the absence and presence of 128 μg/mL fluconazole as described in Methods. Libraries were prepared, sequenced, and mapped, and then read counts within the coding sequences of each gene were tabulated and averaged across the duplicates. Each dot represents one annotated gene. Key genes required for innate resistance to fluconazole are highlighted (blue). Mitochondrial genes that cause significant fluconazole resistance when disrupted are indicated (red). Other genes mentioned in the text are highlighted. (B) A portion of the TCA cycle from *S. cerevisiae* and hypothetical inhibition of fluconazole resistance proteins by alpha-ketoglutarate accumulation, which could occur in *kgd1-* and *kgd2-*insertion mutants but not *idh1-* and *idh2-*insertion mutants. (C) Growth of *idh2Δ pdr1Δ* mutants, single mutants, and wild-type parental strain was measured after 20 hr incubation in SCD medium at 30°C following a 2000-fold dilution from stationary phase pre-cultures. Each data point indicates the average of 6 technical replicates and smooth curves illustrate the average IC50, slope, and maxima obtained from non-linear regression using the sigmoid equation.

Resistance of *C. glabrata* to fluconazole is conferred, in part, by a drug efflux pump (Cdr1), homologs of PI transfer protein (Pdr16, Pdr17), and a drug-responsive transcription factor (Pdr1) that induces their expression (40–44). We observed *CDR1*, *PDR16*, *PDR17*, and *PDR1* among the strongest outliers below the main diagonal (Fig. 4A, blue squares) with z-scores of −4.4, −10, −3.1, and −2.7, respectively. We also observed *UPC2A* at 18 standard deviations below the main diagonal (z-score = −18). Upc2A confers fluconazole resistance by inducing expression of the direct target of the drug (Erg11) as well as other enzymes that function in the ergosterol biosynthetic pathway (45, 46). *DAP1*, encoding a heme-binding protein important for Erg11 function and fluconazole resistance (47), also exhibited fluconazole hypersensitivity (z-score = −4.5). In *S. cerevisiae*, Dap1 physically interacts with the product of *YHR045w* (48), and *yhr045wΔ* mutants correlated strongly with *dap1Δ* mutants (Pearson correlation of 0.86 (49)) in terms of their chemical interaction profiles (50). Insertions in *YHR045w* gene of *C. glabrata* (CAGL0J00297g) also resulted in hypersensitivity to fluconazole (z-score = −3.6). These findings demonstrate that genes conferring innate fluconazole resistance in *C. glabrata* can be identified readily through insertion profiling.

Another 17 genes were found to promote innate resistance to fluconazole. The *STV1* gene appeared below the main diagonal (z-score of −10.1), in agreement with a previous report of fluconazole hypersensitivity when *stv1* is disrupted with *Tn7* transposon (41). Stv1 is a component of the Golgi-localized V-ATPase complex, while a related protein Vph1 is a component of the vacuole-localized V-ATPase complex (51). Transposon insertions in *VPH1* caused significant resistance to fluconazole (z-score of +3.4). As mentioned earlier, most other V-ATPase subunits are not duplicated, are shared by both complexes, and score as essential in the enrichment conditions. Interestingly, a V-ATPase inhibitor caused hypersensitivity to fluconazole in *C. glabrata* (52) similar to the *stv1* insertion mutants. The most significant component revealed by GO term analysis of this group involved *KGD1* and *KGD2* (z-score = −5.7, −7.0), whose products in *S. cerevisiae* form alpha-ketoglutarate dehydrogenase complex that synthesizes succinyl-CoA in the TCA cycle (53, 54). The hypersensitive phenotypes of insertions in *KGD1* and *KGD2* is surprising because the opposite behavior was observed for insertions in *IDH1* and *IDH2* (z-score = +8.4 and +5.4) whose products synthesize alpha-ketoglutarate (see Fig. 4B). These findings suggest that the product of Idh1-Idh2 (alpha-ketoglutarate) may somehow lower intrinsic resistance to fluconazole.

If alpha-ketoglutarate diminishes fluconazole resistance by lowering the function of Pdr1, an *idh2Δ* knockout mutation should cause resistance when introduced into wild-type cells but not when introduced into *pdr1Δ* mutant cells. To test this possibility, we knocked out *IDH2* in BG14 (wild-type) and CGM1094 (*pdr1Δ* derivative of BG14 (55)) strains and quantified growth at different levels of fluconazole (Fig. 4C). The *idh2Δ* single mutant was 2.81-fold more resistant to fluconazole (IC50 = 66.7 ± 6.3 μg/mL) than wild-type (IC50 = 23.7 ± 1.6 μg/mL) and the difference was highly significant (p = 1.4E-5, Student’s t-test). Importantly, the *idh2Δ pdr1Δ* double mutant and the *pdr1Δ* single mutant exhibited equal sensitivity to fluconazole (IC50 = 3.6 ± 0.1 and 3.4 ± 0.1 μg/mL, respectively). These findings suggest that natural biosynthesis of alpha-ketoglutarate in mitochondria can suppress the functions of Pdr1 (modeled in Fig. 4B).

In addition to *IDH1*, *IDH2*, and *VPH1*, 197 other genes above the main diagonal were significant (z-score >= 3). The striking majority of this group (135 genes) encode proteins required for mitochondrial functions such as translation, respiration, TCA cycle (excepting *KGD1* and *KGD2*), and lipid biosynthesis (Fig. 4, red circles). Previous studies showed mutations disrupting *PGS1*, *SUV3*, and *MRPL4* genes, which are critical for diverse mitochondrial functions, also confer resistance to fluconazole, likely through hyperactivation of Pdr1 (41, 56). These three genes (z-scores = 13.9, 7.0, and 4.3, respectively) fall within the minor diagonal formed by the other mitochondrial genes. Previous studies showed that complete loss of the mitochondrial chromosome causes activation of Pdr1, up-regulation of Cdr1 and Pdr16, and resistance to fluconazole (44, 57). By generating new pools of transposon insertions in *pdr1Δ* knockout mutants, *rho0* mutants, and other backgrounds, it should be possible to disentangle the regulatory networks that converge on Pdr1 and to identify novel Pdr1-independent regulators of fluconazole susceptibility (58).

Numerous additional genes contributed significantly to fluconazole susceptibility. Out of 65 non-mitochondrial genes that are significantly resistant to fluconazole when disrupted by transposons, the top three genes encode subunits of the SKI complex (*SKI2, SKI3*, and *SKI8*; z-scores = 12.8, 9.5, and 11.4), which recruits the exosome and facilitates 3’ to 5’ degradation of mRNAs (59). We were unable to find any precedence in the literature for involvement of the SKI complex in susceptibility to fluconazole in any species. GO term analysis of the remainder of this group yielded only one significant process: Glycosphingolipid Biosynthesis (GO:0006688, P-value = 4.7E-4), which contained catalytic and regulatory subunits of MIPC synthase (encoded by *CSG2*, *SUR1*).

An additional gene of interest that lies below the diagonal is *CNA1* (z-score of −3.0; Fig. 4), which encodes the catalytic subunit of the calcium-activated protein phosphatase known as calcineurin (60). The *CNB1* gene, which encodes the regulatory subunit of calcineurin, contained no insertions in the starting pool probably because it is very small (528 bp) and distant from the centromere (911 kb). Calcineurin plays a minor role in resistance to fluconazole and other azoles (11, 60). However, calcineurin plays a major role in the promotion of cell survival during exposure to high fluconazole in *C. glabrata* (41, 60, 61) and many other yeasts including *S. cerevisiae* (62) and *C. albicans* (61, 63). This calcineurin-dependent “tolerance” to azoles can be blocked by calcineurin inhibitors, such as FK506 and cyclosporine, thus converting fluconazole from a fungistat to a fungicide (64). Because the culture conditions involved high doses and long exposure times, we expect calcineurin-deficient mutants to exhibit significant loss of viability during the exposure to fluconazole and thus lower regrowth upon re-culturing in fresh medium lacking fluconazole. The hypersensitive mutants lacking Stv1 or Pdr1 did not exhibit significant cell death in response to fluconazole (41), suggesting these proteins promote resistance mechanisms and not tolerance mechanisms. On the other hand, calcineurin promotes resistance of *C. glabrata* to the echinocandin-class antifungal caspofungin by inducing expression of the drug targets (60, 65). Calcineurin also promotes cell survival in serum and virulence in mouse models of disseminated candidiasis and ocular infection (66). *Hermes* insertion profiling in *C. glabrata* may shed new light on the mechanisms that regulate all these processes.

## DISCUSSION

Efficient and comprehensive methods of functional genomics are critical to understand the principles and molecular mechanisms that distinguish pathogens such as *C. glabrata* from model organisms such as *S. cerevisiae*. Here we develop the *Hermes* transposon from housefly as a powerful new tool for functional genomics research in *C. glabrata*. Unlike CRISPR/Cas9 methods, where mutations are user-guided and not directly sequenced, *Hermes* insertions *in vivo* are nearly random, inexpensive to generate, and easy to profile directly, thus probing the functions of all regions of the genome without guidance. In spite of modest bias of *Hermes* insertions toward nTnnnnAn sequences and nucleosome-free regions, all annotated genes of *C. glabrata* and *S. cerevisiae* can be disrupted if the pool size is large enough to achieve sufficient coverage. Interestingly, while transposons tend to reinsert close to the site from which they were launched, we observed a significant difference between *C. glabrata* and *S. cerevisiae* in the degree of this form of bias. By launching *Hermes* from a centromere-containing plasmid, which would be clustered in the nucleus close to all the other chromosomal centromeres, we observed a much stronger and lengthier bias of insertions near centromeres of *C. glabrata* relative to *S. cerevisiae.* This difference probably reflects some previously unappreciated difference in chromosome architecture or compaction, which potentially could be visualized using chromosome conformation capture and 3D representation (67).

Since their last common ancestor, *C. glabrata* has lost hundreds more ancestral genes than *S. cerevisiae* (8), all of which are non-essential in *S. cerevisiae* except for *BIR1*. *BIR1* is likely annotated incorrectly as a pseudogene in *C. glabrata* because the in-frame stop codon, a sequence conforming to a programmed +1 ribosomal frameshift, and the essential C-terminal domain are strongly conserved in all six species of the *Nakaseomyces* clade. Programmed +1 ribosomal frameshifting occurs broadly in eukaryotes and is utilized to regulate the expression of antizyme (Oaz1 in yeasts), thus coupling polyamine biosynthesis to cellular concentration of spermidine (68). While more research will be needed to establish the regulation of *BIR1* expression through +1 ribosomal frameshifting, the implications in *C. glabrata* are noteworthy. *BIR1* overexpression has been shown to increase virulence of *Aspergillus fumigatus* in mouse models of lung infection (69), possibly even when truncated to lack the essential C-terminal domain. Overexpression of only the N-terminal portion of Bir1 accelerates the growth rate of wild-type *S. cerevisiae* in culture (70). Perhaps more importantly, Bir1 assembles via its C-terminal domain into the chromosomal passenger complex, which is crucial for proper segregation of chromosomes in mitosis from yeast to human (35). Thus, the regulated expression of Bir1 with or without its C-terminal domain could potentially alter virulence traits or the rate of chromosomal aneuploidy (71), which have been previously associated with acquisition of fluconazole resistance (5) and host infection by *C. glabrata* (6). Deficiency of *BIR1* function in *S. cerevisiae* also causes synthetic lethal interactions with several different kinetochore genes, perhaps explaining why these same genes have become essential in *C. glabrata* but not *S. cerevisiae*.

With the assistance of machine learning (21), the essentialome of *C. glabrata* has been defined for the first time. While several kinetochore genes appeared to be essential in *C. glabrata* and not *S. cerevisiae*, 15 genes essential genes in *S. cerevisiae* involved in mRNA splicing were scored as non-essential in *C. glabrata*. This could reflect the diminished number of introns and intron-containing genes in *C. glabrata* relative to *S. cerevisiae* (30, 37). However, other explanations cannot be ruled out without validation and further testing. In the 1-to-1 orthology group, 84% of essential genes in *S. cerevisiae* were also scored by machine learning as essential in *C. glabrata*. Even so, false positives occurred due to repetitive sequences and small gene sizes while false negatives may also occur if essential genes are located in regions of the genome that are sparsely inserted with transposons or if the essential gene products contain large non-essential domains at their 3’ ends (e.g. *PAN1*). Calling of essential genes from transposon insertion datasets could be improved by also incorporating datasets from diploids, where very few heterozygous gene knockouts exhibit strong fitness defects (13, 21).

Until now, methods for quantifying the complexity of the sequencing libraries and normalizing different experiments have been rudimentary. To improve the situation, we developed a midLC statistic that empirically quantifies the complexity of a library independent of jackpots (insertions that occur at early phases of culture and are over-represented) and the degree of over-sequencing (see Materials and Methods). The degree of library over-sequencing can be quantified relative to the midLC and then used to normalize datasets from different runs prior to analyses. Accurate measures of library complexity and jackpot frequency can help define bottlenecks in the protocols that would lower complexity and diminish the usable capacity of sequencing runs.

The power of transposon mutagenesis and insertion profiling was also illustrated through the identification of non-essential genes that become essential in other conditions. Original efforts made use of the bacterial *Tn7* transposon that was first inserted into purified genomic DNA *in vitro* and then recombined into the *C. glabrata* genome (41). Individual clones were then screened manually for resistance and hypersensitivity to fluconazole (41) and caspofungin (72). Now, much larger pools of insertion mutants can be generated and screened by combining *in vivo* transposition with deep sequencing technology. With this approach, we confirm large contributions of Upc2A, Pdr1, Pdr16, Pdr17, and Cdr1 to the innate fluconazole resistance of *C. glabrata* through their ability to increase fluconazole efflux or fluconazole target expression. We also report dozens of other genes whose products contribute to fluconazole resistance (e.g. Stv1, Kgd1-2 complex) and hypersensitivity (e.g. Ski2-3-8 complex, mitochondrial functions including Idh1-2). Our findings suggest that alpha-ketoglutarate biosynthesis by Idh1-2 in mitochondria naturally diminishes Pdr1 function in fluconazole resistance. Though these experiments with gene knockout mutants validate our findings with transposon insertions, more direct experiments will be necessary to determine how alpha-ketoglutarate (or some related metabolite) interacts with Pdr1. The conditions employed herein only revealed the genes with rather strong contributions to fluconazole susceptibility. With modifications to the experimental design, more genes with more subtle contributions will surely become apparent. By iterating the process in mutants lacking Pdr1 (or others), networks of gene function can be inferred.

The pools of *Hermes-NATr* insertions described here can immediately be utilized for exploring *C. glabrata* susceptibility to any number of clinical and experimental antifungals. Additionally, new pools can be generated in the extremely diverse sub-clades that define the *C. glabrata* species group (73), enabling exploration of their vast phenotypic diversity. The pCU-MET3-Hermes launchpad developed here may even be useful for pool generation in other species of the *Nakaseomyces* clade, thereby enhancing our understanding of genomic and phenotypic plasticity that underlays the evolution of virulence in this group of emerging pathogens (74).

## MATERIALS AND METHODS

### Plasmids, organisms, and culture conditions

The plasmid pCU-MET3-Hermes was constructed by double digestion of pCU-MET3 (22) with SpeI and XhoI and 3-way ligation with a 2.1 kb SpeI-NotI and a 2.3 kb NotI-XhoI fragment from pSG36 (17). DNA sequencing confirmed that *Hermes* transposase coding sequences were placed downstream of the methionine-repressible MET3 promoter of *C. glabrata* on the centromeric plasmid bearing the *Hermes-NATr* transposon. This plasmid was introduced into strain BG14, a *ura3-* derivative of *C. glabrata* BG2 (23), by transformation using the lithium acetate protocol (23). Individual transformants were purified and stored frozen at −80°C until use.

To generate pools of *Hermes-NATr* insertion mutants, single colonies of BG14 [pCU-MET3-Hermes] were inoculated into 100 mL synthetic SCD medium lacking uracil, cysteine, and methionine and shaken for 3 days at 30°C in either a single 500 mL glass culture flask (pools 1 and 2) or forty 16 × 150 mm glass culture tubes which were then pooled (pool 3). The cells were then pelleted, resuspended in 600 mL SCD medium containing 0.1 mg/mL nourseothricin and 1 mg/mL 5-fluoroorotic acid and shaken overnight at 30°C in 2 L culture flasks. This step was repeated once more. Finally, 60 mL of these partially enriched cultures were pelleted and resuspended in 600 mL of the same medium and cultured as before. The highly enriched cultures were pelleted, resuspended in 15% glycerol, and frozen in aliquots at −80°C. Aliquots were thawed and sequenced according to the QIseq protocol below or diluted 10-fold into 10 mL fresh SCD medium, grown overnight at 30°C, and diluted 100-fold into 300 mL fresh SCD medium containing or lacking 128 μg/mL fluconazole. These cultures were shaken at 30°C for 24 hr, then the cells were pelleted, washed 1x in SCD medium to remove residual drugs, resuspended in 300 mL fresh SCD medium, and cultured at 30°C for an additional 2 days. Cells were then pelleted, resuspended in 15% glycerol, and frozen in aliquots at −80°C as before.

The *IDH2* gene (CAGL0I07227g) was knocked out in strains BG14 and CGM1094, an isogenic derivative of BG14 that carries *pdr1Δ* mutation (55), using a variation of the standard protocol (11). Briefly, 5’ and 3’ homology arms located upstream and downstream of the *IDH2* gene were PCR amplified from BH14 genomic DNA, fused to an intervening PCR product of *URA3* from *S. cerevisiae*, and transformed into BG14. The Ura+ colonies were screened by PCR using primers listed in Supplemental Table S3 to identify mutants where *IDH2* was replaced with *idh2Δ∷URA3*. A verified *idh2Δ* mutant (named AGY07), a verified *idh2Δ pdr1Δ* double mutant (AGY04), and the parent strains were grown to saturation in SCD medium, diluted 2000-fold into fresh medium containing variable concentrations of fluconazole in 96-well dishes, mixed, and incubated at 30°C for 20 hr. Optical density at 650 nm was then measured for 6 technical replicates. IC50 was calculated independently for each replicate by fitting the data to a standard 3-parameter sigmoid equation using non-linear regression (Kaleidagraph v4.5 software).

### Genomic DNA extraction, QIseq, data processing and visualization

To extract genomic DNA, 100 mg of thawed cell pellets were washed three times in 1 mL deionized water and extracted using Quick-DNA Fungal/Bacterial Miniprep kit (Zymo Research). A total of 2.4 μg of purified gDNA was fragmented by sonication to average size of ~350 bp in four separate aliquots using a Diagenode Picoruptor. The fragmented DNA was then end repaired, ligated to splinkerette adapters (Supplemental Table S3), size selected with AMPure xp beads, and PCR amplified in separate reactions using transposon-specific and adapter-specific primers as detailed previously (15). Samples were then PCR amplified twice (Supplemental Table S3) to enrich for insertion sites and to attach Illumina P5 and P7 adapters that were not indexed (pool Cg-1) or indexed (pools Cg-2, Cg-3). PCR products were purified with AMPure XP beads (Beckman Coulter Life Sciences), mixed with phiX-174, loaded into MiSeq instrument (Illumina) and 75 bp of each end was sequenced using custom primers specific for Hermes right inverted repeat and P7 (Supplemental Table S3). Detailed protocols and primer sequences are available upon request. DNA sequence reads (Cg-2 and Cg-3) were de-multiplexed using CutAdapt (75) mapped to the *C. glabrata* CBS138 reference genome (version 32) using Bowtie2 (76), and any mapped reads with a quality score <= 20 or a mismatch at nucleotide +1 were removed to eliminate ambiguous mappings. The number of DNA sequence reads that map to each unique site in the genome was tabulated using a custom script in Python. Data from multiple sequencing runs were visualized using the IGV genome browser (77) after scaling the number of reads at each site according to the following formula: reads × 20 + 100 (16).

### Computational methods

The libraries prepared from 3 different pools were sequenced to different levels (ranging from 7.2 to 21.5 million mapped reads), and therefore required normalization before they can be combined and analyzed. For normalization, we developed a back-sampling approach that simultaneously estimates the frequency of jackpots (the frequency of reads that occur at one or more over-represented sites) and midLC (mid-library complexity, or the number of reads that produce 2x midLC unique insertion sites after eliminating jackpots). Briefly, the list of mapped reads was sampled randomly at multiple different depths (100, 400, 1600, 64000, etc.) in triplicate up to the maximum and the number of unique sites at each depth was recorded. The frequency of unique sites at each depth (y-axis) was charted against depth (x-axis) and the data were fit to a 3-parameter sigmoid equation y = (1−jackpot) / (1+(x/midLC)^slope) using non-linear regression (Kaleidagraph v4.5 software). The output produced estimates of jackpot frequency (percentage of high frequency sites), mid-library complexity or midLC (after excluding jackpots, the read depth where half map to unique sites and the remainder map to those same sites), and slope factor. In all three libraries, jackpots were low (< 0.04%) while midLC varied over a 1.6-fold range (Supplemental Figure S4). The library derived from pool Cg-1 had the highest number of mapped reads and the lowest midLC, with a ratio between the two values of 259. In contrast, the more complex libraries from pools Cg-2 and Cg-3 were sequenced to 65.1 and 63.3 times the midLC, respectively, or about 4-fold lower depth than Cg-1. For comparison, libraries prepared from *Hermes* insertion pools in haploid and diploid *S. cerevisiae* (21) yielded midLC’s that averaged 1.06- and 1.74-fold higher than *C. glabrata* libraries (Supplemental Figure S4). A compilation of six *mini-Ac/Ds* libraries from *S. cerevisiae* (16) exhibited far higher midLC and jackpots (Supplemental Figure S4). Thus, midLC, jackpots, and depth of sequencing can vary significantly between species, transposons, and methodologies.

To identify essential genes in *C. glabrata*, the datasets obtained from pools Cg-2 and Cg-3 were normalized to correct for 4.0-fold under-sequencing relative to that of Cg-1 and then combined. The combined dataset was analyzed by an 8-feature machine learning algorithm that was previously used to identify essential genes in *C. albicans*, *S. pombe*, and *S. cerevisiae* (21). The features were weighted based on a validated training dataset of 692 essential and 1756 non-essential genes from *S. cerevisiae* (21).

To quantify insertion site sequence bias, raw data from pools 1, 2, and 3 were compiled into a single table and nucleotides at positions +2 and +7 were imported from the reference genome. The read counts at each of the 16 possible sites were summed for each pool and compared to all such sites in the CBS138 reference genome. To quantify centromere bias, the read counts at each insertion site from pool 1 were binned into 1 kb segments from the centromere and averaged across all 26 chromosome arms of *C. glabrata* and all 32 arms of *S. cerevisiae* (21).

To calculate z-scores of each gene, the log(2)ratio of experiment/control was divided by the local standard deviation, which was calculated as follows. The data from two biological replicates of control culture conditions were tabulated for each gene and used to calculate average and log(2)ratio. The table was sorted from highest to lowest average and then the 40-gene running average and running standard deviation of the log(2)ratios were calculated. The running standard deviation (y-axis) was charted against the average (x-axis) and fit to a power function y = m1 + m2*x^m3 using non-linear regression (Kaleidagraph) for all genes with average read counts >= 6. The parameters obtained from the best-fits were then used to calculate local standard deviation of the log(2)ratios for each gene in the experimental and control datasets after slight normalization of the replicates.

Multiple sequence alignment using MUSCLE and phylogenetic tree analysis using PhyML were implemented using SEAVIEW v4 (78).

### Data availability

Raw DNA sequence data were deposited at the Sequence Read Archive (NCBI) with the bioproject ID PRJNA625944. Tables of mapped sequence reads as well as unmappable BG2 and CBS138 sites are available for download from the authors upon request. The .bed files used for IGV genome browser will be available upon request and downloadable from the Candida Genome Database (http://www.candidagenome.org).

## ACKNOWLEDGMENTS

The authors thank Brendan Cormack (Johns Hopkins School of Medicine), Henry Levin (National Institutes of Health, USA), Reed Wickner (National Institutes of Health, USA) and John Adams (University of South Florida) and members of their teams for plasmids, strains, and technical advice on the design and implementation of insertion profiling and QI-seq. We thank Benoît Kornmann and Agnès Michel (University of Oxford) for advice and for mini-Ac/Ds datasets. Zhuwei Xu and Brendan Cormack generously provided genomic DNA sequences of strain BG2 prior to publication. ANG and RMS were supported by National Institute of General Medical Sciences (T32-GM007231). ANG, WT, and KWC were supported by grants from National Institutes of Allergy and Infectious Disease (R21-AI130722) and Johns Hopkins University (Discovery award). AL was supported by a fellowship from the Edmond J. Safra Center for Bioinformatics. RS was supported by the Israel Science Foundation (grants no. 715/18, 757/12). JB was supported by Israel Science Foundation (grant no. 997/18).

## SUPPLEMENTAL FIGURES

**Supplemental Figure S1.**
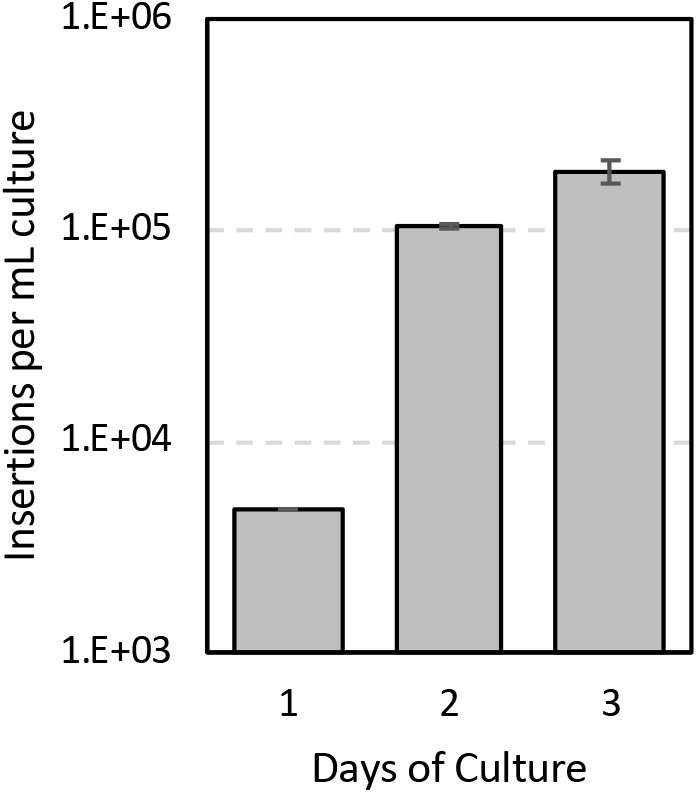
Hermes-NAT1 insertions in *C. glabrata*. Replicate cultures of *C. glabrata* strain BG14 [pCU-MET3-Hermes] were grown in SCD culture medium lacking uracil, cysteine, and methionine to induce expression of transposase and the number of insertion mutants per mL of culture was determined by counting colonies that grew on SCD+NAT+5FOA agar medium after 2 days of incubation at 30°C.

**Supplemental Figure S2.**
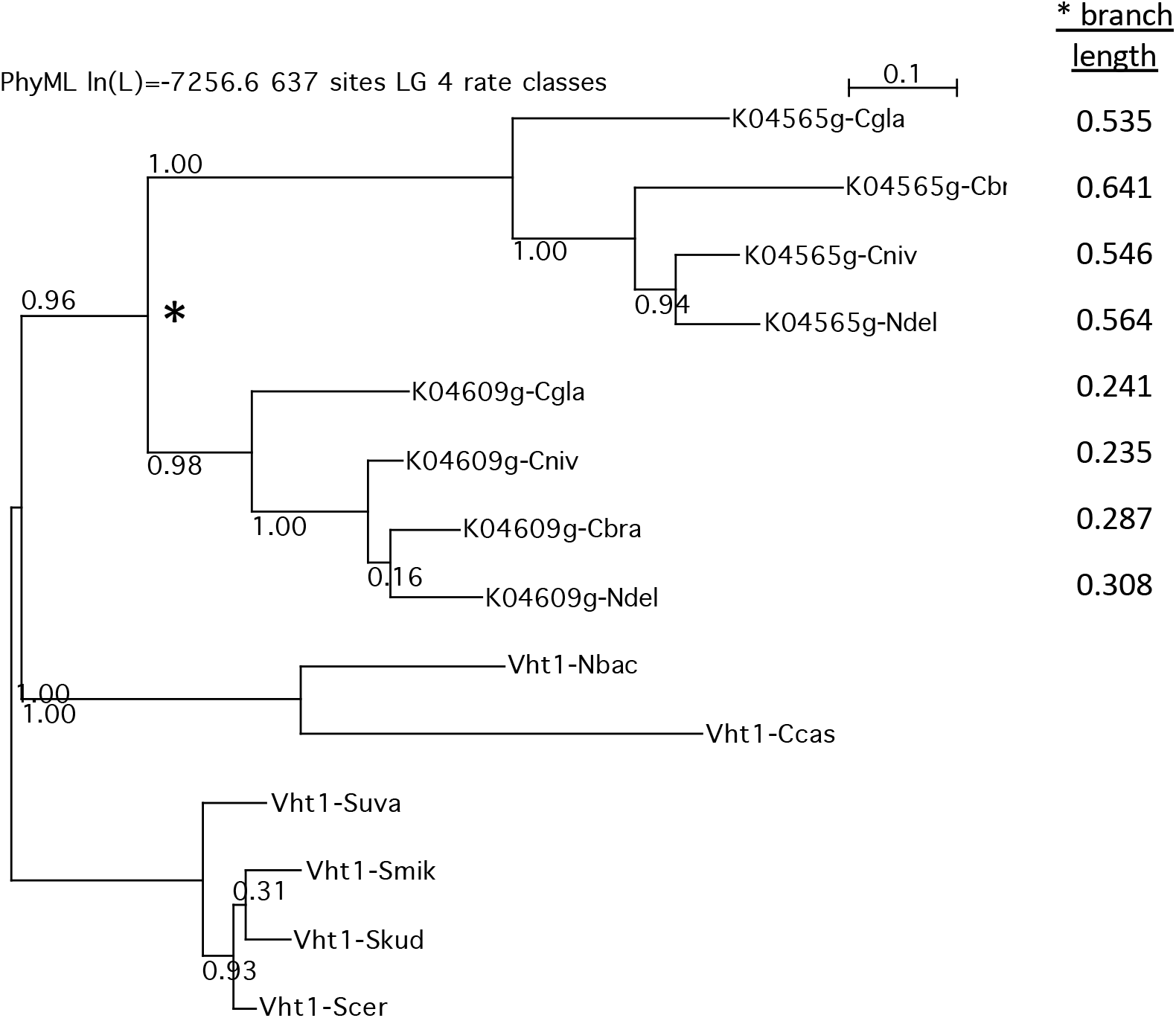
Phylogenetic tree of Vht1-related proteins from *Nakaseomyces* and *Saccharomyces* clades. Protein sequences related to Vht1 of *S. cerevisiae* were identified by BLAST, aligned by MUSCLE, and analyzed by PhyML using Seaview v4.0 software (78). Bootstrap values are indicated at each node and the total lengths each branch relative to the gene duplication event (star) are indicated.

**Supplemental Figure S3.**
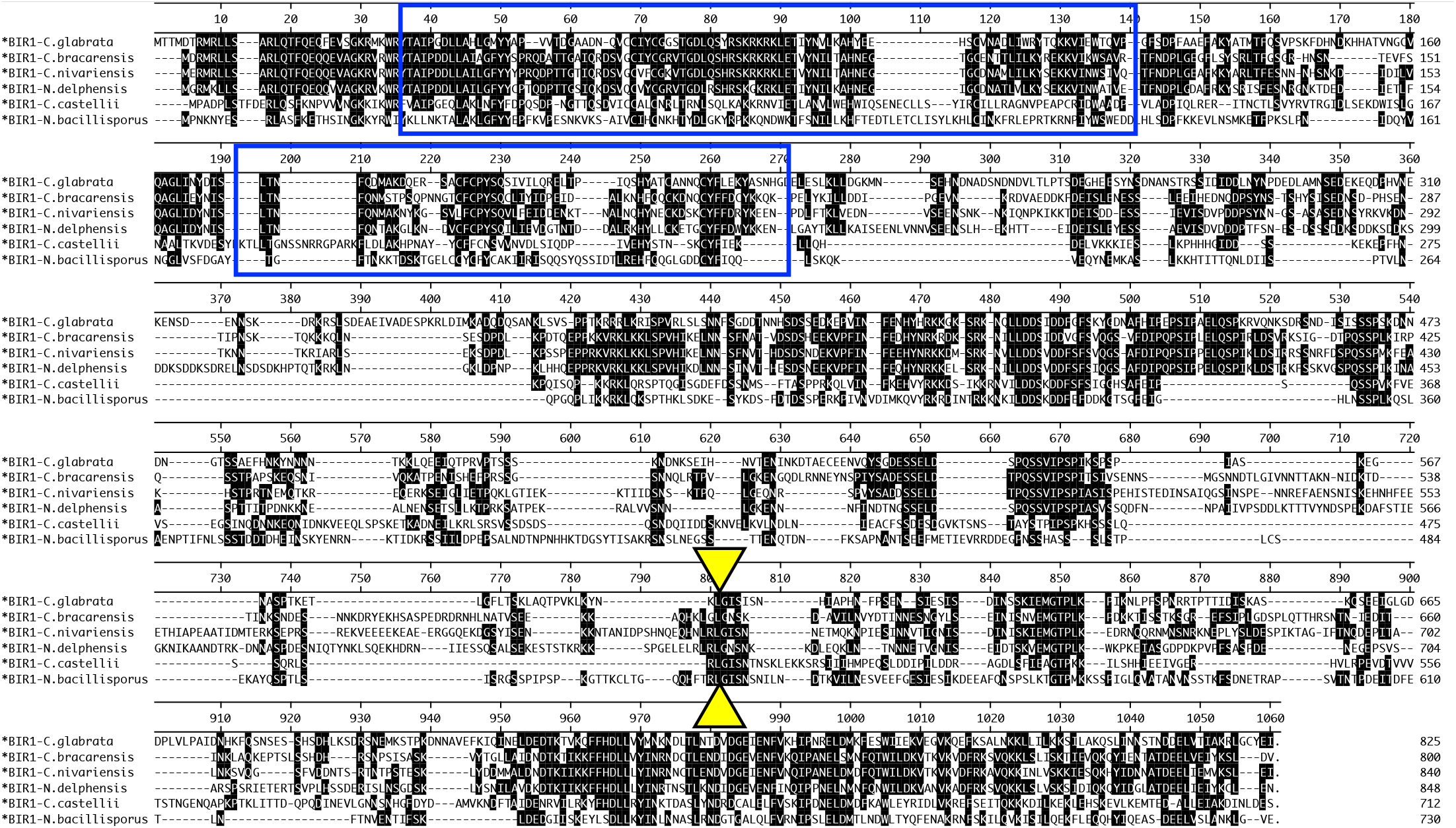
Multiple sequence alignment of Bir1 orthologs from *Nakaseomyces*. DNA sequences from the indicated species were edited to remove the central “A” within the conserved CUU-A-GGC programmed +1 ribosomal frameshift sequence (yellow triangles) and then aligned using MUSCLE. Blue boxes indicate approximate positions of conserved BIR repeats.

**Supplemental Figure S4.**
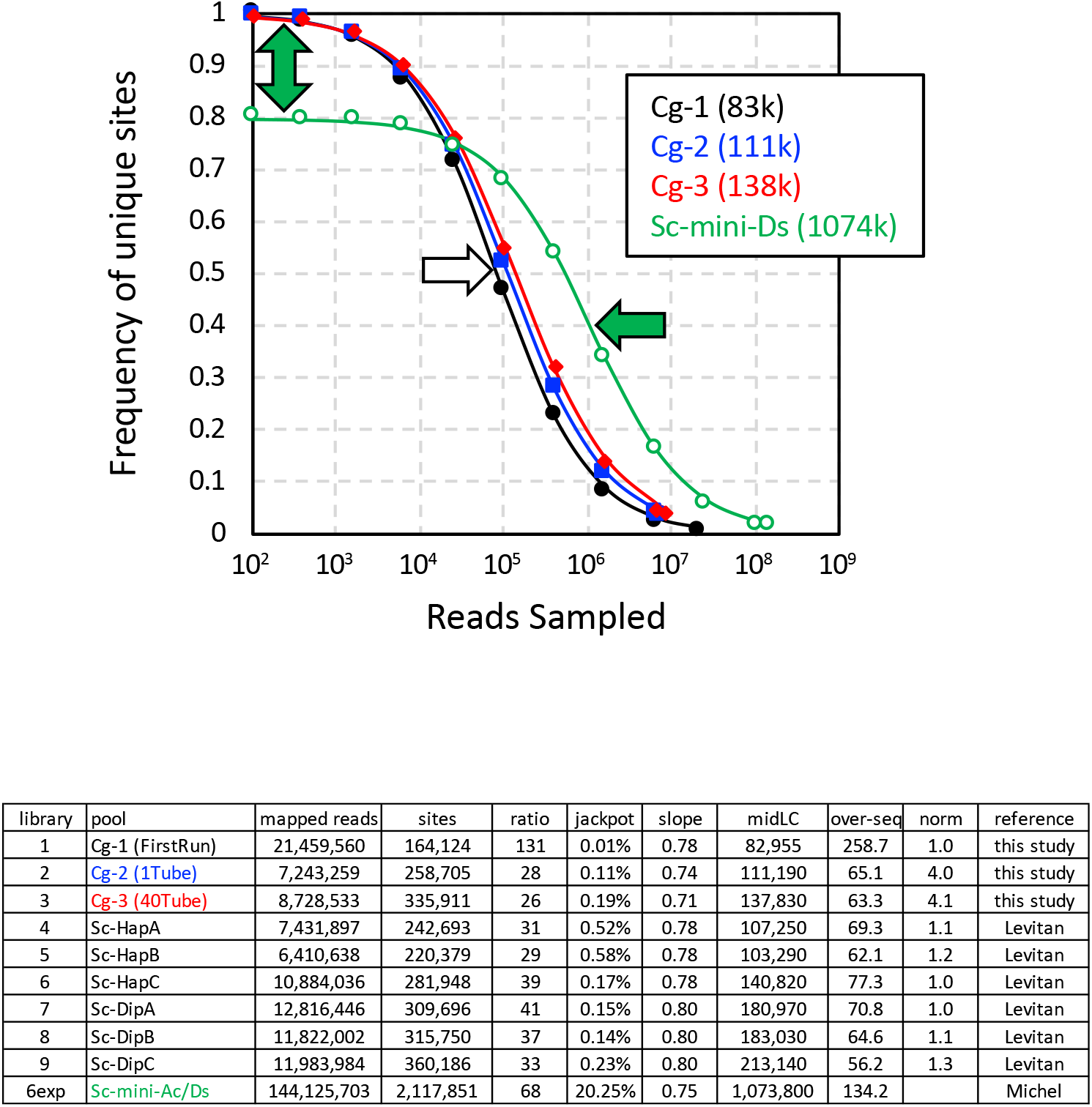
Back-sampling approach for quantification of jackpots, mid-library complexity (midLC), and normalization. Libraries of *Hermes-NAT1* insertions in *C. glabrata* (this study) and *S. cerevisiae* (21) and the sum of six *mini-Ac/Ds* pools in *S. cerevisiae* (16) were sequenced, and mapped. The lists of mapped reads were back-sampled at different depths, and the frequencies of unique sites at each depth were charted in the graphic. Data were fit by non-linear regression to a 3-parameter sigmoid equation (smooth curves) as described in Materials and Methods. White arrow indicates the positions of midLC for all 3 *C. glabrata* libraries. Green arrow indicates the position of midLC from six combined *mini-Ac/Ds* datasets, after excluding 20% jackpots (green double arrow). The table shows values obtained from non-linear regression (jackpot, slope, midLC). The degree of over-sequencing was calculated by dividing the total number of mapped reads by the midLC. This revealed that library Cg-1 was over-sequenced relative to Cg-2 and Cg-3 by about 4-fold, so the latter libraries were normalized accordingly prior to combining for analyses by machine learning. The Sc libraries were sequenced to similar depths and not normalized before combining (21).

## SUPPLEMENTAL TABLES

**Table S1. Essentiality Scores of annotated genes in *C. glabrata* and *S. cerevisiae*.** All annotated genes in *C. glabrata* and *S. cerevisiae* were scored by machine learning for essentiality and arranged by orthology between the two species. The length of each gene and the percentage that is unmappable are also indicated. From the Saccharomyces Genome Database, annotations of essentiality, gene name, and gene description are also included.

**Table S2. Z-scores of annotated genes in *C. glabrata* that alter susceptibility to fluconazole.** For each annotated gene, the total number of reads associated with each *Hermes* insertion site within the coding sequences were tabulated and averaged across replicate experiments after normalization. Z-scores were calculated. Data from earlier studies (11, 41) were also included.

**Table S3. Oligonucleotides utilized in this study.** Adapters and PCR primers used for QIseq were all modified from (15).

